# Visuomotor adaptation modulates the clustering of sleep spindles into trains

**DOI:** 10.1101/2021.10.27.466169

**Authors:** Agustín Solano, Luis A. Riquelme, Daniel Perez-Chada, Valeria Della-Maggiore

## Abstract

Sleep spindles are thought to promote memory consolidation. Recently, we have shown that visuomotor adaptation (VMA) learning increases the density of spindles and promotes the coupling between spindles and slow oscillations, locally, with the level of spindle-SO synchrony predicting overnight memory retention. Yet, growing evidence suggests that the rhythmicity in spindle occurrence may also influence the stabilization of declarative and procedural memories. Here, we examined if VMA learning promotes the temporal organization of sleep spindles into trains. We found that VMA increased the proportion of spindles and spindle-SO couplings in trains. In agreement with our previous work, this modulation was observed over the contralateral hemisphere to the trained hand, and predicted overnight memory retention. Interestingly, spindles grouped in a cluster showed greater amplitude and duration than isolated spindles. The fact that these features increased as a function of train length, provides evidence supporting a biological advantage of this temporal arrangement. Our work opens the possibility that the periodicity of NREM oscillations may be relevant in the stabilization of procedural memories.

**CONTRIBUTION STATEMENT:** Ever since the discovery of memory systems, the study of the mechanisms supporting the consolidation of declarative and procedural memories has progressed somewhat in parallel. We now know, however, that structures originally thought of as purely declarative such as the hippocampus, participate in the consolidation of procedural tasks. Recently, we showed that sleep predicts long-term motor memory through the local synchrony between fast sleep spindles and slow oscillations, a mechanism initially described for the consolidation of declarative memories. Novel evidence has linked the rhythmicity in the occurrence of spindles to memory stabilization. This framework proposes that temporally clustered spindles into *trains* of two or more separated by 3-6 seconds, may favor the reinstatement and subsequent reprocessing of previously acquired memories. This temporal arrangement may facilitate mnemonic replay and neocortical integration. In the present study, we show that motor learning promotes the organization of spindles into trains, locally, over the contralateral hemisphere, and that this modulation predicts overnight memory retention. Spindle grouping also augmented the proportion of spindle-SO couplings in trains. Importantly, spindles in a cluster increased their duration and amplitude as a function of train length, pointing to a physiological benefit of this temporal organization.

## INTRODUCTION

The last decade has seen remarkable progress in the identification of the neural signatures of sleep-dependent consolidation. Overnight reactivation of hippocampal memories appears to depend on sleep spindles, slow oscillations (SO) and their precise synchrony (Maingret et al., 2016; Ladenbauer et al., 2017; Helfrich et al., 2018; Muehlroth et al., 2019; Navarro-Lobato and Genzel, 2019). Recently, we have shown that motor learning, specifically visuomotor adaptation (VMA), also promotes the coupling between spindles and SOs (Solano et al., 2021). The fact that the level of spindle-SO synchrony predicts overnight memory retention opens the possibility that common mechanisms operate in the stabilization of declarative and motor memories.

Sleep spindles are bursts of oscillatory thalamocortical activity thought to support the reactivation of newly formed memories (Rasch and Born, 2013; Staresina et al., 2015; Schönauer, 2018). Individual spindles may promote memory consolidation through the modulation of intrinsic features such as their amplitude (Bergmann et al., 2012; Barakat et al., 2012; Boutin et al., 2018), duration (Morin et al., 2008; Laventure et al., 2016), power (Ngo and Staresina, 2020) and density (Nishida and Walker, 2007; Barakat et al., 2011; Albouy et al., 2013a; Solano et. al., 2021). In the last few years, it has been argued that the rhythmicity in their occurrence may also be a relevant parameter influencing memory stabilization (Lecci et al., 2017; Antony et al., 2018, Boutin and Doyon, 2020). Specifically, during NREM sleep (stages 2 and 3) spindles tend to cluster into *trains* of two or more, separated by inter-spindle intervals (ISI) of 3 to 6 seconds (Evans and Richardson, 1995; Achermann and Borbély, 1997; Antony et al., 2018). This ISI is believed to reflect a refractory period. It has been proposed that recently acquired memories would be reinstated during a spindle oscillation and subsequently reprocessed during the following refractory period, thereby protecting memory reinstatement from potential interference (Antony et. al., 2018; 2019). Boutin and collaborators (2018a, 2019, 2020) have in fact shown that some metrics of spindle trains, such as the ISI, predict offline gains in motor sequence learning. Yet, whether clustering modulates spindle’s intrinsic features such as duration and/or amplitude remains unknown. This information may be key to unveil the physiological advantage of spindle grouping.

Here, we followed up on our previous findings (Solano et al., 2021), to examine if VMA learning promotes the temporal organization of sleep spindles into trains. We also explored if the spindle-SO coupling which, as we showed is modulated by VMA, is also influenced by spindle rhythmicity. Animal studies have shown that the cluster of spindles into trains may increase the recruitment and synchronization of a greater number of thalamic and cortical neurons involved in spindle generation (Contreras and Steriade, 1996), which from the EEG recording would manifest as an increment in spindle amplitude. Clustering may also strengthen corticothalamic connectivity and long-range synchronization (Bonjean et al., 2011), which would reveal as longer spindle duration. To shed light on the biological relevance of spindle clustering, we also quantified the relationship between their duration/amplitude and train length. We found that VMA increased the proportion of spindles in trains locally, over the contralateral hemisphere to the trained hand. Importantly, clustered spindles were longer and larger than isolated spindles, pointing to a physiological benefit of this temporal organization.

## MATERIALS AND METHODS

### Participants

Ten healthy volunteers (5 females, age: (mean ± SD) 24.3±3.1 years old) completed the whole study. All subjects were right-handed, and none of them declared neurological nor psychiatric disorders. Only subjects fulfilling the criteria for good sleep quality (Buysse et al. 1989; Johns 1991) were included in the study.

All volunteers signed the informed consent approved by the Ethics Committee of the Hospital de Clínicas, University of Buenos Aires (approved on Nov, 24, 2015, and renewed every year), which complies with the Declaration of Helsinki in its latest version, and with the National Law on the Protection of Personal Data.

### Experimental paradigm and procedure

Subjects performed a visuomotor adaptation task (VMA) consisting of moving a cursor that represented their hand, from a start point located in the center of a computer screen to one of 8 visual targets arranged concentrically, using a joystick. The latter was controlled with the thumb and index finger of the right hand, while vision of the hand was occluded. One cycle consisted of eight trials, whereas there were 11 cycles in a block.

Subjects were instructed to perform fast shooting movements through each of the eight targets presented pseudorandomly within a cycle. There were three different types of trials throughout the study. During perturbed trials a clockwise 45-degree visual rotation was imposed to the cursor relative to the movement of the hand (ROT). During null, i.e., unperturbed trials (N) in which no visual rotation was applied, the movement of the cursor directly mapped onto the hand movement. Finally, during error-clamp (EC) trials, visual feedback was manipulated to provide fake “straight” paths to the target that mimicked those generated during correct trials. This was accomplished by projecting the actual movement to the straight line with some additional variability (mean error=0°, SD=10°). EC trials prevent further learning, and allow estimating memory retention based on the internal state of the motor system (Villalta et al., 2015; Criscimagna-Hemminger & Shadmehr, 2008).

### Experimental Design

The experimental design of this study is fully described in a previous publication derived from the same dataset (Solano et al., 2021). Briefly, we conducted a within-subject experiment consisting of three sessions separated by 7 days each: i) a familiarization session, ii) a visuomotor adaptation session (VMA) in which a visual rotation (ROT) was applied, and iii) a control session (CTL) in which subjects performed unperturbed, null trials (N). The order of the VMA and CTL sessions was counterbalanced. In the VMA and CTL conditions, subjects performed the task before and after sleep on Day 1 and Day 2. On Day 1, they performed one block of N trials followed by six blocks of ROT/N trials, depending on the condition. On Day 2, they performed two cycles of EC trials to assess memory retention, followed by 6 blocks of ROT/N trials, depending on the condition, and three blocks of N trials. In each session, subjects went to bed ∼10 min after performing the task on Day 1 for a full night of sleep, and a polysomnographic (PSG) recording was obtained throughout the night.

### EEG recording and processing

For the PSG recording, eleven surface electroencephalogram (EEG) electrodes were placed over prefrontal, motor and parietal areas (FC1, FC2, FC5, FC6, C3, C4, P3, P4), and over the midline (Fz, Cz, Pz). Both mastoids were used as references. Electrooculography (EOG) and electromyography (EMG) signals were also obtained. All signals were acquired at 200 Hz.

EEG, EOG and EMG signals were bandpass-filtered to facilitate sleep scoring (EEG: 0.5-30 Hz; EOG: 0.5-15 Hz; EMG: 20-99 Hz). All PSG recordings were sleep staged manually, according to standard criteria (Iber, 2004). Namely, 30-second epochs were classified as either Wake (W), Non Rapid Eye Movement (NREM1, NREM2 and NREM3), or Rapid Eye Movement (REM) stage.

Sleep spindles (10-16 Hz) and slow oscillations (0.5-1.25 Hz) were automatically identified from the EEG signal corresponding to the stages NREM2 and NREM3, using previously reported algorithms (see below).

#### Sleep spindles detection

The algorithm was based in the work of Ferrarelli and collaborators (2007) and Mölle and collaborators (2011). First, EEG signal for each channel and session was bandpass-filtered (10-16 Hz) before calculating the instantaneous amplitude (IA) and instantaneous frequency (IF) by applying the Hilbert Transform (Tort et al., 2010). The IA was used as a new time series and smoothed with a 350 ms moving average window. Next, those segments that exceeded an upper magnitude threshold (90^th^ percentile of all IA points) were labeled as potential spindles. Spindles onset and offset were defined as the time points in which the signal dropped below a lower threshold (70^th^ percentile of all IA points). Potential spindles with a duration between 0.5 and 3 seconds were labeled as true spindles. Finally, spindles were further classified into two types according to their frequency: slow spindles (<12 Hz) and fast spindles (≥12 Hz) (Mölle et al., 2011; Cox et al., 2017).

#### Detection of Slow oscillations (SO)

The algorithm was based on that reported by Mölle and collaborators (2011) and Antony & Paller (2016). First, the EEG signal was bandpass-filtered (0.5-1.25 Hz). Then, we identified zero crossings and labeled them as positive-to-negative (PN) or negative-to-positive (NP). Those EEG segments between two NP zero crossings were considered slow oscillations if they lasted between 0.8 and 2 seconds. Next, we computed their peak-to-peak (P-P) amplitude. Finally, we determined the median of the P-P amplitudes for each channel, each subject and each session, and kept those SOs with a P-P amplitude greater than the median value (Mizrahi-Kliger et al., 2018).

We quantified a spindle-SO coupling if a spindle had its maximum peak-to-peak amplitude during the course of a SO.

#### Spindles Trains

In a recent publication, Boutin and Doyon (2020) defined a *spindle train* as a group of two or more locally occurring sleep spindles separated by a maximum inter-spindle interval (max_ISI) of 6 seconds. This criterion was based on the distribution of ISIs reported in previous studies (Evans & Richardson, 1995; Achermann and Borbély, 1997; Antony et al., 2018). Spindles occurring at longer ISIs than the max_ISI threshold were thus considered isolated spindles. In this approach the max_ISI is a fixed parameter invariant across channels and subjects. To examine if this criterion adjusted well to observed data, we contrasted it with that obtained from applying an adaptive method used in neurophysiology to identify bursts of action potentials (Kapucu et al., 2012). This approach, which allows to compute a max_ISI to identify spike bursts, is composed of the following steps, illustrated in Supplementary Figure 1. First, ISIs are quantified and the corresponding histogram is constructed. Second, a Cumulative Moving Average (CMA) curve is derived from the histogram according to the function

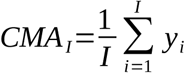

where *y*_*i*_ is the number of observations for each bin of the ISI histogram, *i*=*1*, …,*N*, and *N* the total number of ISI bins. CMA_*I*_ is the *I*th value of the CMA curve, with *I*≤*N*. Then, based on the skewness of the CMA curve, a scaling parameter, *α*, is determined, which will then be used to identify the max_ISI (*a*=*1* if *Skewness* < *1*; *α*=*0*.*7* if *1* ≤ *Skewness* < *4*; *α*=*0*.*5* if *4* ≤ *Skewness* < *9*; and *α*=*0*.*3* if *Skewness* ≥ *9*). The max_ISI then corresponds to the ISI where the CMA curve falls from its maximum (*CMA*_*max*_) to *α*CMA*_*max*_. We applied this method on data from each channel and each subject, and then computed the median of the max_ISI for each sleep stage (NREM2 and NREM3) and each experimental session. This yielded four max_ISI values per subject. To optimize the ability of the algorithm to resolve one train from another, we kept the minimum max_ISI out of the four values.

### Data analysis

#### Behavior

Motor performance was measured based on the angle defined by the movement direction of the joystick and the line segment connecting the start point and target position (pointing angle). Trial-by-trial data were converted into cycle-by-cycle time series by computing the median pointing angle for each cycle of eight trials and each subject. To assess memory retention for each subject the pointing angle of the two EC cycles was expressed as a percentage of the pointing angle asymptote (median of the last block of learning), and then averaged.

#### EEG signal

##### SO and Spindle measures

Given that some electrodes came off after the first two hours of EEG recording, we focused the EEG analysis on the first sleep cycle.

The following measures were computed: spindle duration (msec), peak-to-peak amplitude (μV), inter-spindle interval (ISI), proportion of spindles in trains (number of spindles in trains/total number of spindles) and the proportion of spindle-SOs in trains (number of spindle-SO in trains/total number of spindle-SO).

To assess how these measures differed between VMA and CTL sessions we computed their relative difference according to the function ((VMA-CTL)/CTL*100) for each EEG channel and each subject (Huber et al., 2004, 2006; Solano et al., 2021). To illustrate the spatial distribution of the effects we report the results in topographic maps (MNE-Python; Gramfort et al., 2013). Furthermore, given that our previous work has shown that VMA modulates NREM oscillations contralaterally to the trained hand (Solano et al., 2021), we contrasted the data pooled across the electrodes of the left hemisphere (LH: FC1, FC5, C3 & P3) against the data pooled across the electrodes of the right hemisphere (RH: FC2, FC6, C4 & P4). As the midline (Fz, Cz, Pz) may capture electrical activity from both hemispheres it was excluded from this analysis.

To investigate whether the intrinsic features of spindles were influenced by being temporally organized into trains, we examined the amplitude and duration of spindles in trains versus isolated spindles. We chose these measures because they reflect the size of neural populations active during a spindle and their level of synchronization, which are often modulated by learning (Morin et al., 2008; Bergmann et al., 2012; Barakat et al., 2012; Boutin et al., 2018; Laventure et al., 2016). Moreover, to compare trains of different length, we focused on the features of the first and last spindle in a train. We hypothesized that if being part of a train was somewhat advantageous, then these features would be potentiated in clustered vs isolated spindles, and this potentiation would become stronger as a function of train length. To this aim, we first calculated the median duration and amplitude of isolated spindles and that of the first and last spindle in a train, for each EEG channel and each subject, and then computed the mean duration and amplitude across channels for each subject. Finally, for each initial and final spindle in a train, we computed the percent difference in amplitude and duration relative to that of isolated spindles, and statistically contrasted them.

### Statistical analysis

Statistical analyses were carried out using R (version 3.4.1; R Core Team, 2017). Statistical differences were assessed at the 95% level of confidence, and were carried out by fitting Linear Mixed Models (LMM, using the ‘lmer’ function implemented in the ‘lme4’ package in R, Bates et al., 2015). Random intercepts and random slopes of LMMs were estimated for each subject to take into account the repeated measures across sleep stages and hemispheres. Data from each of the four electrodes that were pooled together for each hemisphere were considered replicates. The response variable was the relative difference between sessions ((VMA-CTL)/CTL*100). The fixed effects were the sleep stage (NREM2 and NREM3), the cerebral hemisphere (Left and Right) or the type of spindle (fast and slow), depending on the analysis. To assess the statistical significance of fixed effects and obtain p-values, we used F tests or t tests with Kenward-Roger’s approximation of the degrees of freedom (Halekoh and Højsgaard, 2014). Note that in cases where the value of a replicate deviates beyond 2.5 median absolute deviation (MAD, Leys et al., 2013), the degrees of freedom may not result in an integer.

A one sample t-test was used to compare the max_ISIs found by applying the CMA algorithm with the fixed 6 seconds threshold.

Finally, to statistically assess whether train length impacts on the duration and amplitude of spindles, we first conducted repeated-measure correlations using the “rmcorr” package in R (Bakdash and Marusich, 2017). The repeated-measure correlation coefficient, r_RM_, represents the strength of the linear association but accounting for the repeated observations for each subject. Next, we ran LMMs to assess whether the intrinsic features of spindles varied along the train, by comparing the first and last spindles. The response variable was the relative difference of the intrinsic features for the first and last spindle in a train relative to isolated spindles. The fixed effects were the position of the spindle (first or last) and the length of the train. The latter was included in the model as a covariate of no interest.

## RESULTS

### Organization of sleep spindles into trains

After identifying individual sleep spindles, we computed the ISIs to explore their periodicity. Figure 1.A shows the distribution of ISIs during NREM2 and NREM3. As previously shown, the distribution of ISIs is right-skewed and reveals that most spindles are separated by ∼3-6 second intervals. Based on this observation, Boutin and Doyon (2020) set a max_ISI of 6 seconds to determine if a group of spindles constitute a train. We compared this criterion to that obtained from applying the adaptive algorithm created by Kapucu and collaborators (2012). This approach yielded a max_ISI of 5.16 ± 0.5 seconds (mean ± SE). Given that the two criteria did not statistically differ (one sample t-test, t(9)=-1.7, p=0.12), we used the fixed 6 second criterion for train quantification.

**Figure 1.**
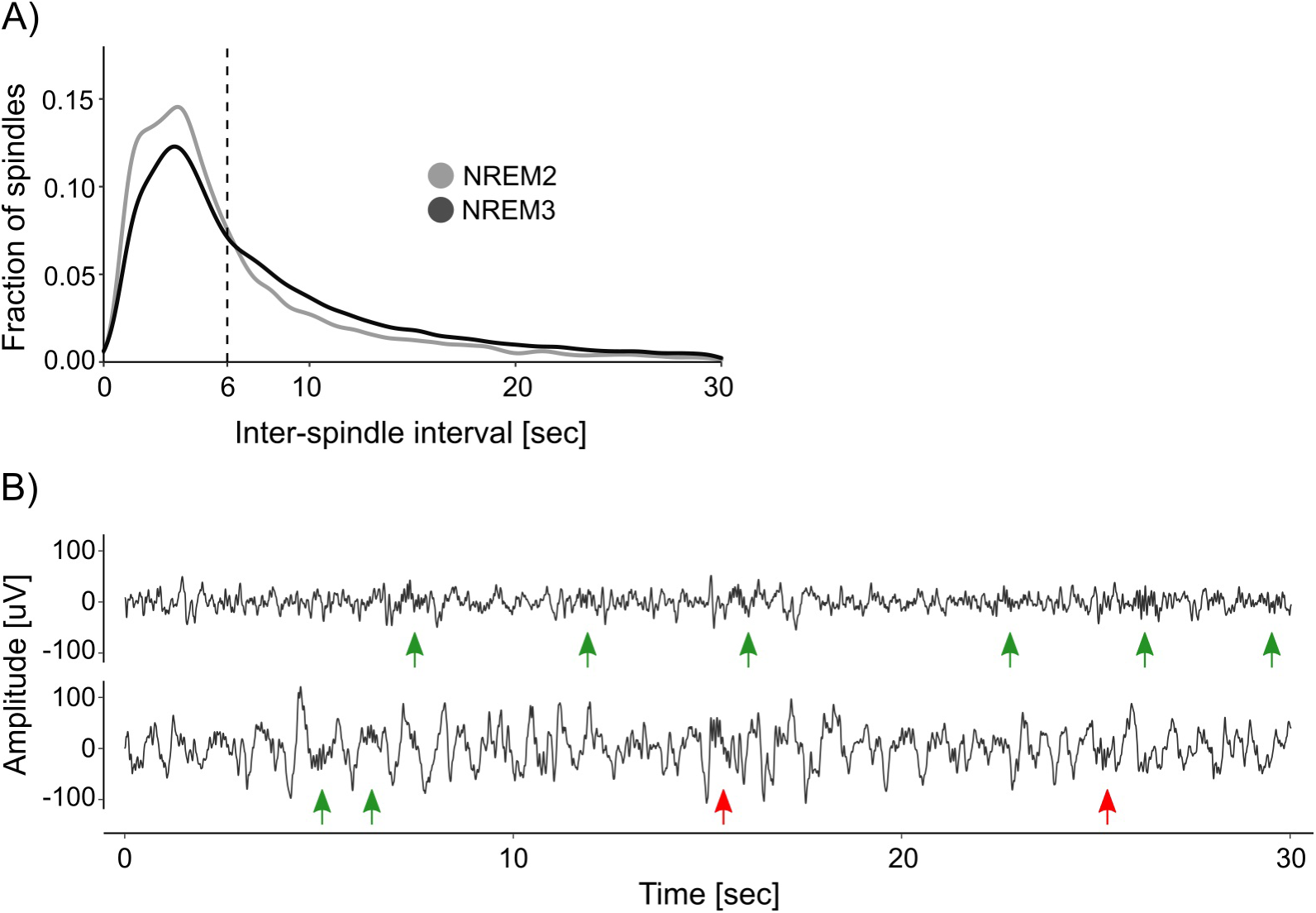
Organization of sleep spindles into trains. *A) Distribution of inter-spindle intervals*. Shown is the distribution of inter-spindle intervals (ISIs) during NREM2 and NREM3. *B) Examples of spindle trains*. Two 30 second segments of sleep EEG are depicted, one from NREM2 (top trace) and one from NREM3 (bottom trace). Green arrows indicate sleep spindles clustered into trains, whereas red arrows indicate isolated spindles. Note that according to the 6 second max_ISI criterion, there are two trains of three spindles on the top trace and one train of two spindles on the bottom trace.

Figure 1.B depicts two EEG traces from NREM2 and NREM3 that illustrate spindle trains (green arrows) and isolated spindles (red arrows). As detailed in Table 1, most trains contained between 2 and 3 spindles, whereas few trains may encompass 7 spindles or more.

**Table 1.** Proportion of trains as a function of the number of spindles. Shown is the total amount, proportion and cumulative proportion of trains according to their length during NREM2 and NREM3.

| # Spindles<br>in train | Total amount |  | Proportion (%) |  | Cumulative proportion (%) |  |
| --- | --- | --- | --- | --- | --- | --- |
|  | NREM2 | NREM3 | NREM2 | NREM3 | NREM2 | NREM3 |
| 2 | 2246 | 5655 | 40 | 48 | 40 | 47 |
| 3 | 1109 | 2774 | 20 | 23 | 60 | 71 |
| 4 | 705 | 1420 | 13 | 12 | 73 | 83 |
| 5 | 437 | 759 | 8 | 6 | 81 | 89 |
| 6 | 304 | 443 | 5 | 4 | 86 | 93 |
| 7 | 208 | 240 | 4 | 2 | 90 | 95 |
| 8 | 150 | 190 | 3 | 2 | 93 | 97 |
| 9 | 94 | 112 | 2 | 1 | 95 | 98 |
| 10 | 78 | 74 | 1 | 1 | 96 | 99 |
| 11 | 65 | 41 | 1 | 1 | 97 | 99 |

### Visuomotor adaptation modulates the clustering of spindles into trains

To explore if the temporal clustering of spindles into trains was modulated by motor learning, we computed the proportion of spindles in trains relative to the total number of spindles during NREM2 and NREM3, for each EEG channel and each subject. Figure 2.A depicts the topographic maps for NREM2 (top row) and NREM3 (bottom row). We found a learning-related increase in the proportion of spindles organized into trains over the contralateral hemisphere to the trained hand, specifically during NREM3 (relative change in proportion, mean ±SE: NREM2: LH=-1.3±2.1 %, RH=-0.3±1.9 %; NREM3: LH=5.7±1.9 %, RH=-0.2±1.3; LMM stats, sleep stage by hemisphere interaction: F(1,121.9)=4.59, p=0.03; t-test for Left hemisphere during NREM3 vs zero: t(8.99)=2.14, p=0.03). In consistency with our previous work (Solano et al., 2021), we found that VMA increased the proportion of spindles in trains differentially for fast (≥12Hz) but not slow (<12 Hz) spindles during NREM3 (relative change in proportion, mean ±SE: fast spindles=8.5±2.1 %, slow spindles=-3.4±0.8 %; LMM stats, main effect of spindle type: F(1,8.9)=5.21, p=0.04).

**Figure 2.**
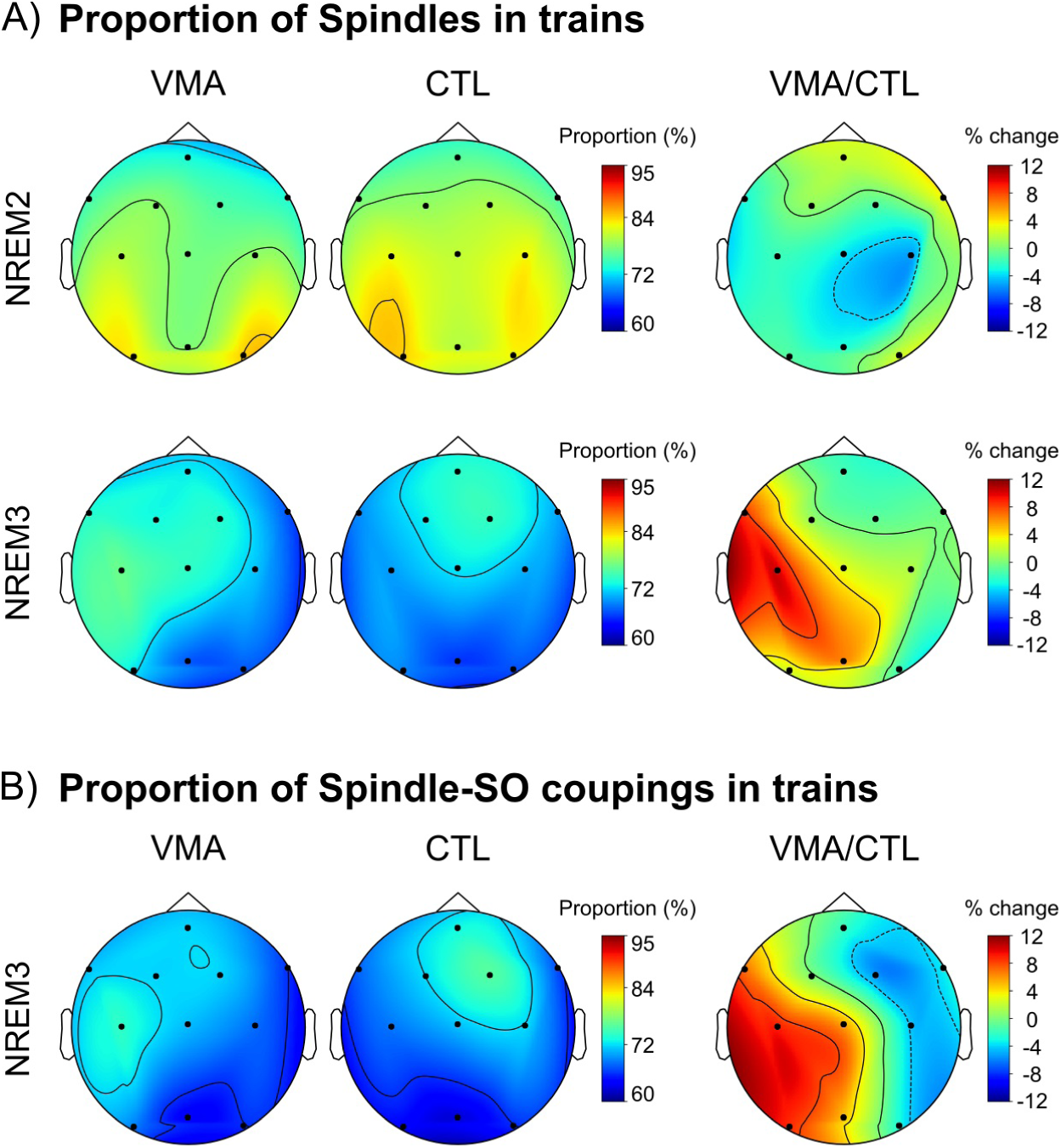
Local modulation of the proportion of spindles and spindle-SO couplings clustered into trains. *A) Visuomotor adaptation increases the proportion of spindles in trains*. Shown are the topographic plots for the proportion of spindles in trains (number of spindles in trains relative to the total number of spindles) corresponding to the VMA session, the CTL session and their relative difference (VMA/CTL) computed according to the function ((VMA-CTL)/CTL*100), during NREM2 (top row) and NREM3 (bottom row). *B)* V*isuomotor adaptation increases the proportion of spindle-SO couplings clustered into trains*. Shown are the topographic plots for the proportion of spindle-SO couplings in trains (number of spindle-SO couplings in trains relative to the total number of spindle-SO couplings) corresponding to the VMA session, the CTL session and their relative difference (VMA/CTL), during NREM3. Colorbars represent the proportion for the VMA and CTL conditions, and their percent change, respectively.

In order to examine whether this interhemispheric modulation was associated with long-term memory, we next correlated the interhemispheric difference in the proportion of spindles in trains with overnight memory retention. We found a significant correlation (Pearson correlation, r=0.68, p=0.045), suggesting that the temporal organization of spindles into trains may benefit motor memory stabilization during sleep.

Lastly, given that in our previous work we showed that VMA augmented the density of slow oscillations coupled with spindles, we examined if it also increased the proportion of spindle-SO couplings clustered into trains. As shown in Figure 2.B, we found a learning-related interhemispheric modulation for this measure, driven by the contralateral hemisphere to the trained hand (LMM stats, main effect of hemisphere; F(1,8.95)=8.95, p=0.01).

### Train length modulates intrinsic parameters of the constituting spindles

So far, we have shown that VMA modulates the amount of spindles into trains, locally. But what may be the physiological benefit of this temporal organization? Sleep spindles may provide a time window favoring the reinstatement of recently acquired memories (Buzsaki, 2015; Antony et al., 2019), and thus, memory stabilization (Antony et al., 2018; Boutin and Doyon, 2020). Then, one possibility is that spindle grouping upregulates intrinsic features of the spindles, such as amplitude and/or duration, thereby increasing the instances of memory reinstatement.

To investigate whether the intrinsic features of spindles were influenced by being temporally organized in trains, we explored the percent difference in duration and amplitude between clustered spindles and isolated spindles (the raw values for these intrinsic features are depicted in Supplementary Table 1). Trains with 2 to 6 spindles were considered in the analysis to include 90% of the trains identified for each subject. Figure 3 illustrates the relative differences in duration (Figure 3.A) and amplitude (Figure 3.B), as a function of train length. The plots reveal that both intrinsic features increased as a function of train length, and reached greater values than those of isolated spindles. This positive relationship was statistically confirmed through repeated measures correlations, and held both for the first and last spindle in a train (repeated measures correlations (r_RM_), Duration: First vs. Isolated spindle: r_RM_=0.67, p<0.001; Last vs. Isolated Spindle: r_RM_=0.75, p<0.001; Amplitude: First vs. Isolated spindle: r_RM_=0.58, p<0.001; Last vs. Isolated Spindle: r_RM_=0.48, p<0.001). To explore further whether spindle amplitude and duration varied along the train, we next compared the first and last spindles. A significant effect would speak in favour of an advantage of being part of a cluster. We found that spindle duration and amplitude were larger in magnitude for the last spindle than for the first spindle in a train (LMM stats, main effect of spindle position: Duration: F(1,9)=7.31, p=0.024; Amplitude: F(1,9)=5.31; p=0.046). These results open the possibility that the increment in the proportion of spindles induced by VMA learning may have impacted on long-term memory through the modulation of spindle duration and/or amplitude.

**Figure 3.**
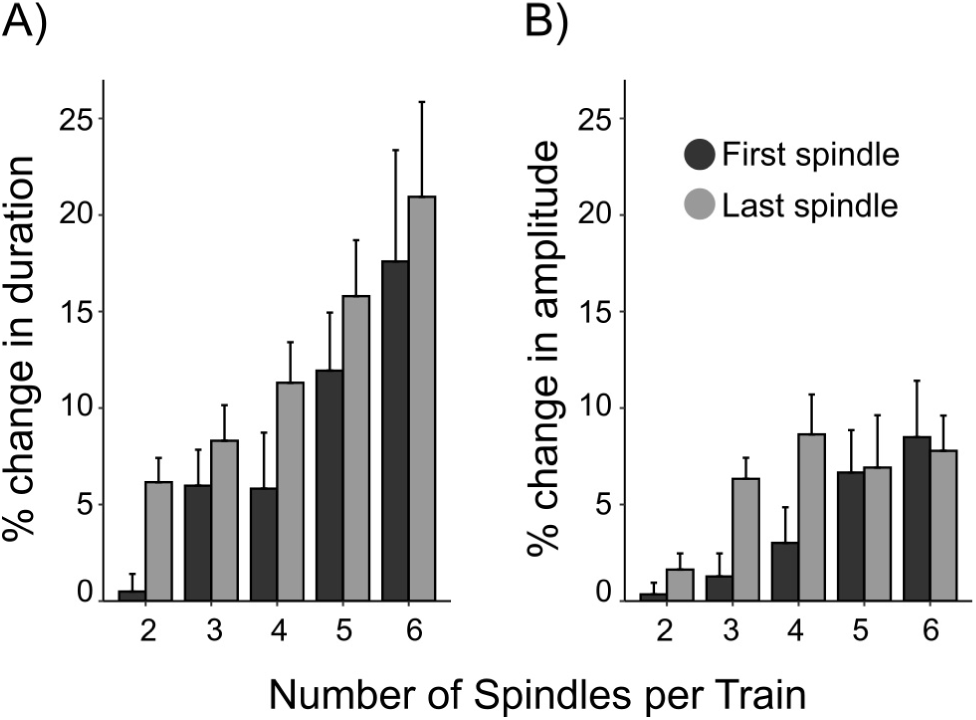
The temporal clustering of spindles modulates their intrinsic features. Barplots represent the mean ± SE of the percent difference in duration (*A*) and amplitude (*B*), for the first and last spindle in a train relative to isolated spindles. This was computed for 90% of trains containing between 2 and 6 spindles. Note that the last spindle in a train tends to be longer and larger than the first spindle (p<0.05).

## Discussion

Uncovering the neural signatures of procedural memory stabilization during sleep is of major relevance to neuroscience research. Recently we found that visuomotor adaptation learning is associated with an increment in the level of coupling between spindles and SOs locally, which predicts overnight memory retention (Solano et al., 2021). Here, we explored further whether VMA also promotes the temporal organization of sleep spindles into trains. We found that VMA increased the proportion of spindles in trains over the contralateral hemisphere of the trained hand; this interhemispheric modulation predicted overnight memory retention. Spindle clustering mostly involved fast spindles during NREM3 -not NREM2. In line with our previous findings, VMA also increased the proportion of spindle-SO couplings organized into trains. The fact that intrinsic features of spindles such as their duration and amplitude augmented with train length points to a physiological advantage of this temporal organization.

Spindle trains are hypothesized to influence memory processing through the alternation between instances of memory reinstatement and refractory periods (Antony et al., 2018, Antony et al., 2019; Boutin and Doyon, 2020). To date, only a few studies have examined the relationship between spindle clustering and memory stabilization. On one hand, Lecci et al. (2017) and Antony et al. (2018) have shown that cluster periodicity (∼0.02 Hz) influences the recall of declarative memory during NREM2 and NREM3. On the other hand, Boutin and collaborators (2018a, 2019) found that motor sequence learning (MSL) reduces the spindle ISI during NREM2, pointing to an increment in the level of clustering, and that this modulation predicted offline gains overnight. Our work contributes further to the understanding of the temporal arrangement of spindles in several ways. First, we show that a type of motor learning heavily dependent on an implicit component increased the clustering of both uncoupled and SO-coupled spindles. Second, this modulation took place specifically during NREM3. Third, the proportion of grouped spindles was augmented over the contralateral hemisphere, suggesting that the network active during learning may reactivate during sleep (Della-Maggiore et al., 2017; Klinzing et al., 2019). Finally, the interhemispheric modulation of spindle clustering predicted overnight memory retention, suggesting a behavioral relevance of this temporal arrangement. Contrary to some of the literature described above, train modulation was specifically observed during NREM3. This may respond to mechanistic differences inherent of the type of learning (declarative vs non-declarative and VMA vs MSL; Krakauer et al. 2019), and/or the experimental design. Further work will be needed to systematically compare tasks.

Although trains may have a role in mnemonic processes, the physiological mechanisms through which spindle grouping modulates memory stabilization remain unexplored. Here we found that both spindle duration and amplitude increased in temporally clustered spindles as a function of train length. Longer spindles may reflect higher cortico-thalamic connectivity and coordination (Bonjean et al., 2011), whereas larger spindles may reflect a greater recruitment of thalamo-cortical neurons (Contreras and Steriade, 1996). We speculate that VMA may impact on long-term memory by promoting the development of longer and larger spindles, thereby facilitating cortico-thalamic synchronicity. Our work points to a physiological advantage in the temporal arrangement of spindles into clusters.

## Supplementary Figure

**Supplementary Figure 1.**
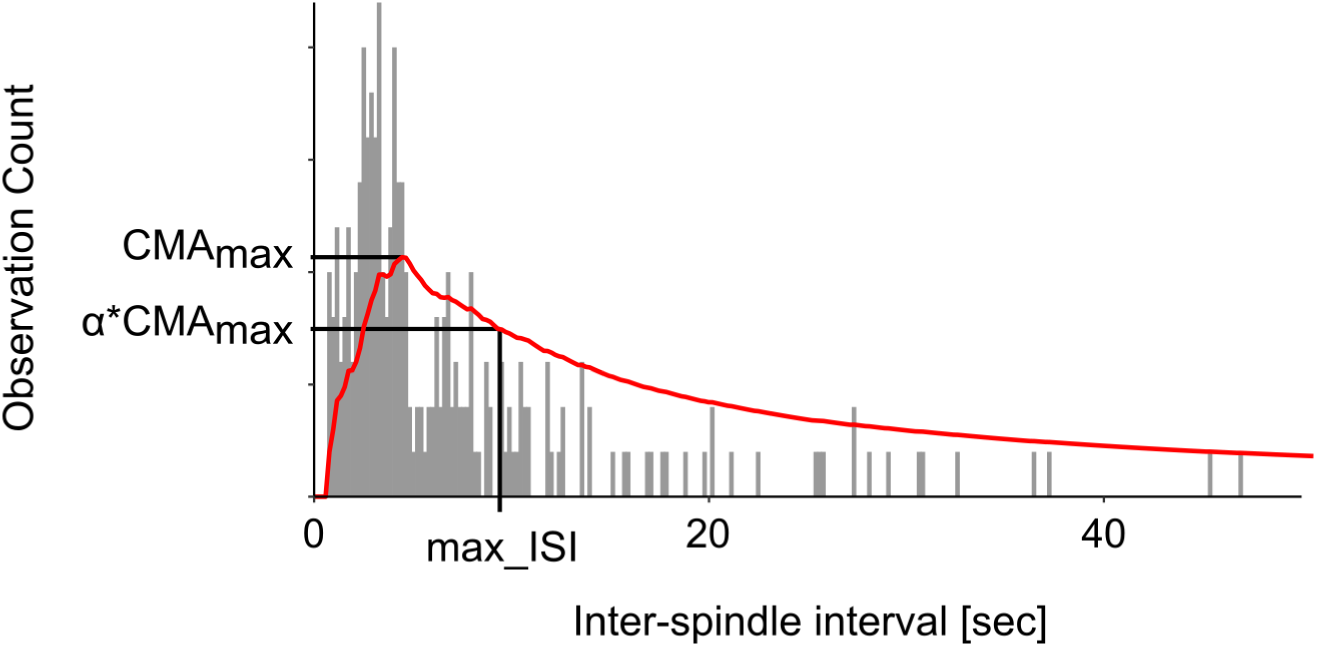
Identification of the max_ISI threshold using the CMA curve. Shown is the histogram of the inter-spindle intervals (ISI) for an example subject (gray bars), and the Cumulative Moving Average (CMA) curve computed for this histogram (red curve). Based on the skewness of the CMA curve, a parameter a is determined and used to scale the maximum of the CMA curve (CMA_max_). The CMA_max_ and the CMA_max_ multiplied by a (a*CMA_max_) are highlighted on the CMA curve. Note that the max_ISI corresponds to the ISI where the CMA curve falls to a*CMA_max_. This max_ISI can be used as the threshold for spindle trains detection.

## Supplementary Table

**Supplementary Table 1.** Shown are the mean and standard error (SE) corresponding to the duration and amplitude of isolated spindles and that of the first and last spindles in a train.

| Number of<br>spindles in<br>Train | Duration (ms) |  |  |  |  |  | Amplitude (uV) |  |  |  |  |  |
| --- | --- | --- | --- | --- | --- | --- | --- | --- | --- | --- | --- | --- |
|  | Isolated<br>Spindle |  | First Spindle |  | Last Spindle |  | Isolated<br>Spindle |  | First<br>Spindle |  | Last<br>Spindle |  |
|  | Mean | SE | Mean | SE | Mean | SE | Mean | SE | Mean | SE | Mean | SE |
| Isolated | 887.0 | 33.5 |  |  |  |  | 52.2 | 3.9 |  |  |  |  |
| 2 |  |  | 890.2 | 31.4 | 940.9 | 35.1 |  |  | 52.5 | 4.1 | 53.1 | 4.0 |
| 3 |  |  | 937.6 | 31.4 | 959.6 | 36.3 |  |  | 52.9 | 4.1 | 55.7 | 4.5 |
| 4 |  |  | 932.2 | 22.9 | 985.3 | 35.6 |  |  | 54.1 | 4.8 | 56.6 | 4.0 |
| 5 |  |  | 987.1 | 26.9 | 1025.7 | 42.0 |  |  | 55.9 | 4.9 | 56.3 | 5.4 |
| 6 |  |  | 1030.2 | 28.9 | 1061.2 | 23.8 |  |  | 56.1 | 3.6 | 55.9 | 3.5 |

